# The transition to postpartum lactation establishes a state of maternal antiviral resilience

**DOI:** 10.64898/2026.10.02.756285

**Authors:** Ariana D. Campbell, Taylor Miller-Ensminger, Montserrat C. Arreguin, Elizabeth A. Kennedy, Peter Hewins, Caroline Stickler, Naomi Esterowitz, Madeleine Assad, Kasia Kready, Pamela A. Silver, Scott Hensley, Kellie A. Jurado

## Abstract

Influenza A virus (IAV) causes severe disease during late pregnancy, yet how maternal adaptations in host defense shape susceptibility following childbirth remains unknown. Here, we show that the postpartum period represents a state of enhanced maternal antiviral immunity. Across postpartum stages, dams infected before weaning exhibited increased survival and reduced disease following lethal IAV challenge compared with nonpregnant females. Postpartum protection was associated with enhanced viral clearance and increased pulmonary antigen-specific CD8^+^ T cells and IL-17A-producing gamma-delta T cells. We identify the transition into lactation as a critical determinant of this protection. Protection extended to highly pathogenic avian influenza. Together, these findings reveal that maternal antiviral immunity remains actively shaped after childbirth, establishing the postpartum lactational period as a distinct state of maternal antiviral resilience.

## Main Text

Maternal physiology undergoes profound adaptations across pregnancy to support fetal development while maintaining maternal health (*1–3*). These adaptations reshape host defense, contributing to heightened vulnerability to severe viral disease during pregnancy in both humans and mouse models (*4–5*). This is particularly evident with influenza A virus (IAV), for which disease severity increases across gestation and peaks during later pregnancy (*6–8*). Despite extensive study of maternal immunity during pregnancy, how antiviral host defense evolves following childbirth remains a critical gap in knowledge.

The postpartum period, or “fourth trimester”, represents a biologically distinct yet understudied phase of maternal physiology (*9*). Rather than simply returning to a pre-pregnancy state, the maternal body undergoes extensive remodeling to support recovery, lactation, and offspring care (*10–14*). However, the accompanying maternal immune adaptations remain poorly defined. Emerging clinical evidence suggests that maternal immunity remains highly dynamic during this transition (*15–17*). Immune-mediated diseases that improve during pregnancy, including rheumatoid arthritis, frequently relapse following childbirth, while altered inflammatory responses, including increased circulating IL-17, have been associated with postpartum complications (*18–21*). Together, these observations suggest that maternal immunity continues to adapt following parturition. Here, we investigated how postpartum adaptations in host defense shape susceptibility to IAV across postpartum stages.

### Early and mid-postpartum mice exhibit enhanced resilience to IAV infection

To define IAV susceptibility across the postpartum period, we developed a mouse model of infection at three distinct postpartum stages. Adult females were intranasally inoculated with a lethal dose of the IAV H1N1 strain PR8 at 1, 7, and 21 days postpartum (early, mid, and late postpartum, respectively) with age-matched nulliparous females (females that have never given birth) serving as controls (Fig. 1A). To standardize caregiving load, litters were culled to six pups on postnatal day 3. Pups were weaned prior to PR8 infection at 21 days postpartum, such that the late postpartum group was evaluated entirely post-weaning. Morbidity and mortality were monitored daily by changes in body mass, core body temperature, and clinical signs. Strikingly, mice infected during the early and mid-postpartum periods exhibited significantly improved survival compared with nulliparous controls (Fig. 1B-C). These dams also better maintained their body mass and core body temperature and exhibited fewer clinical signs prior to and during peak disease (5- and 8-days post-infection (DPI), respectively) (Fig. 1B-C). This enhanced protection also extended to sublethal PR8 infection, with early postpartum dams exhibiting reduced morbidity compared with nulliparous females (fig. S1A). Conversely, dams infected during the late postpartum period succumbed to infection and exhibited survival and disease outcomes comparable to those of nulliparous females (Fig. 1D). These data demonstrate enhanced resilience to severe viral disease during the early and mid-postpartum period that wanes following weaning. Given the robust protection observed during early postpartum, we focused subsequent studies on this stage to define the mechanisms underlying maternal immune resilience.

**Fig. 1.**
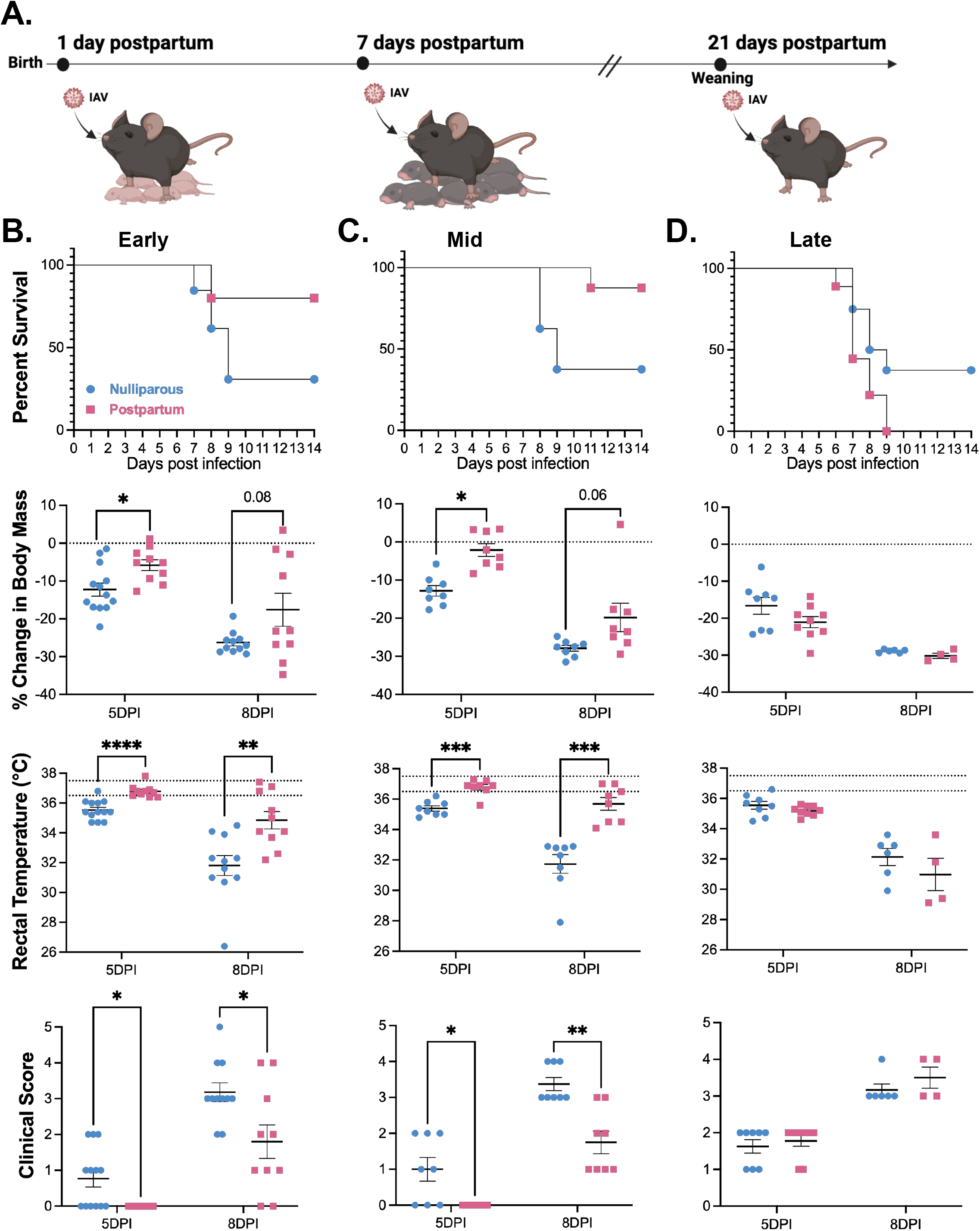
Early and mid-postpartum mice exhibit enhanced resilience to IAV infection. **(A)** Schematic of the experimental timeline. Postpartum dams were intranasally inoculated with 250 TCID_50_ (lethal dose) of influenza A/PR/8/34 (PR8) at 1-, 7-, and 21-days post-birth and compared to age-matched nulliparous females. From top to bottom, mice were monitored daily for survival, changes in starting body mass, rectal temperature, and clinical score for mice infected at the early (1-day postpartum) (n = 10-13/group) **(B)**, mid (7-days postpartum) (n = 8/group) **(C)**, and late (n = 8-9/group) (21-days postpartum) **(D)** postpartum periods. Percent change in body mass, rectal temperature, and clinical score shown at 5- and 8-days post infection (DPI) (corresponding to the onset of disease and peak disease, respectively). For percent change in body mass, dashed lines at 0 indicate starting body mass. For rectal temperature, dashed lines indicate standard core body temperature of laboratory mice (36.5 to 37.5 °C). Error bars represent the mean ± SEM from 3-4 independent experiments with individual mice indicated by symbols. Statistical differences in survival curves were determined by Mantel-Cox test. (B) survival of dams infected at 1-day postpartum, P value = 0.0308, (C) survival of dams infected at 7-days postpartum, P value = 0.0299, (D) survival of dams infected at 21-days postpartum, P value = 0.0578. Statistical differences in changes in body mass and rectal temperature were determined by multiple unpaired t tests and differences in clinical scores were determined by Mann-Whitney tests, all with Holm-Šidák correction for multiple comparisons. *\*P < 0*.*05*, ^*\*\**^*P < 0*.*01*, ^*\*\*\**^*P < 0*.*001*, ^*\*\*\*\**^*P < 0*.*0001*.

A key feature of the postpartum period involves postnatal caregiving (*22*). To determine whether viral infection disrupts postnatal caregiving, we assessed maternal behavior using a pup retrieval assay (*23–24*) at 5- and 8-DPI (fig. S1B). As expected, early postpartum dams retrieved pups significantly faster than nulliparous females with no prior pup exposure, validating robust maternal behavior in postpartum dams (fig. S1, C-D). Importantly, pup retrieval times did not differ between mock-infected and sub-lethal PR8-infected early postpartum dams (fig. S1C). Similarly, early postpartum dams infected with a lethal PR8 dose maintained pup retrieval behavior despite increased disease severity (fig. S1D). Together, these findings demonstrate that postpartum dams maintain caregiving throughout infection, including during peak disease.

### Postpartum dams show distinct immune-mediated mechanisms during IAV infection

Increased morbidity and mortality during IAV infection can result from dysregulated immune responses or impaired viral control (*25*). To determine whether early postpartum protection was associated with differences in viral burden, we quantified infectious virus in the lungs of nulliparous and postpartum mice at 3-, 6-, and 8-DPI following lethal PR8 infection. Consistent with IAV infection kinetics, pulmonary viral burden was highest at 3-DPI and declined over the course of infection (Fig. 2A). No significant differences in viral burden were observed between nulliparous and postpartum dams at 3- or 6-DPI. Strikingly, by 8-DPI, postpartum dams exhibited significantly lower pulmonary viral burdens than nulliparous females, indicating enhanced viral clearance (Fig. 2A).

**Fig. 2.**
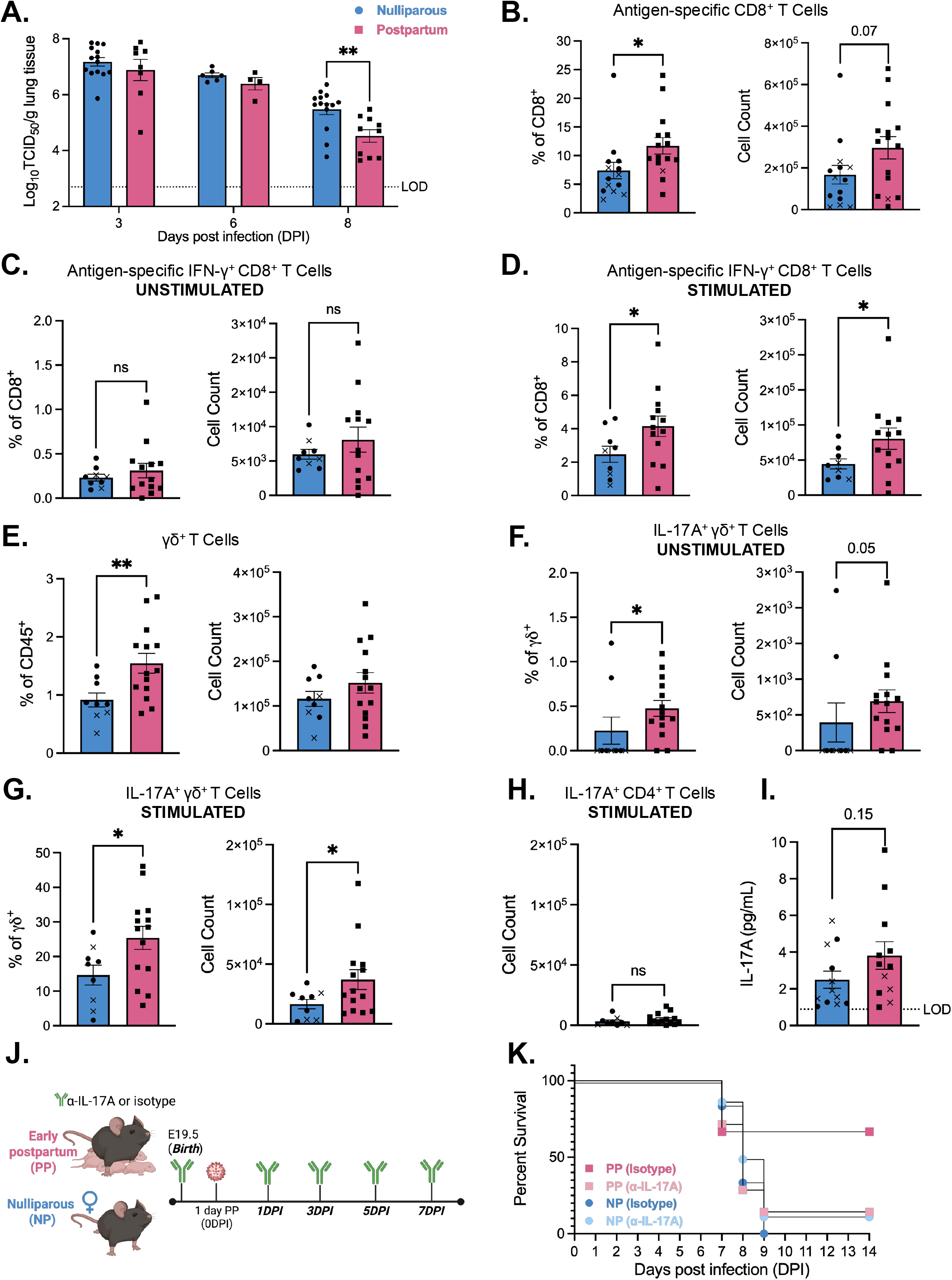
Postpartum dams show distinct immune-mediated mechanisms during IAV infection. **(A)** Nulliparous and 1-day postpartum mice were intranasally inoculated with a lethal dose of PR8 virus. Subsets of mice were euthanized at 3-, 6-, and 8-days post-infection (DPI) to measure viral titers in right lung tissue homogenates via TCID_50_ assay (n = 4-14/DPI and group). **(B-H)** Flow cytometry was performed using cells isolated from whole lung at 8-DPI to assess immune cell type frequencies and counts. **(B)** Frequency and counts of CD8^+^ PR8 nucleoprotein (NP)366-374-specific T cells are shown, followed by frequencies and counts of IFN-γ producing (NP)366-374 CD8^+^ T cells under both unstimulated **(C)**, and stimulated with PMA/ionomycin **(D)** conditions (n = 9-15/group). Frequency and counts of γδ^+^ T cells are shown **(E)**, with **(F)** IL-17A-producing γδ T cells under unstimulated conditions, and **(G)** IL-17A-producing γδ T cells and **(H)** IL-17A-producing CD4^+^ T cells following stimulation with recombinant IL-1β and IL-23 (n = 9-14/group). **(I)** Nulliparous and postpartum dams were euthanized at 8DPI, and left lung homogenates were analyzed for the concentration of IL-17A via LEGENDplex assay (n = 12/group). **(J)** Schematic representation of IL-17A depletion in nulliparous (NP) and postpartum (PP) dams. Mice were administered intraperitoneal injections (i.p.) of anti-IL-17A or isotype control antibody on the day of birth followed by intranasal inoculation of a lethal dose of PR8 virus at 1-day postpartum, with subsequent antibody administrations every other day through 7-DPI. **(K)** Survival of postpartum (PP) and nulliparous (NP) females treated with anti-IL-17A and isotype antibody (n = 3-8/group). Dashed lines indicate the limit of detection (LOD) for each assay. Error bars represent the mean ± SEM from 3-4 independent experiments with individual mice indicated by symbols and “x’s” representing moribund mice. *\*P* < 0.05, ^*\*\**^*P* < 0.01, ns *P* > 0.05. Data were analyzed using two-way ANOVA with Tukey’s multiple comparisons test (A), Unpaired Welch’s t test (B-E and G-I) Mann-Whitney tests (F), and Mantel-Cox test for survival analysis, P value = 0.4123 for all groups, P value = 0.2407 for PP (isotype) versus PP (α-IL-17A) (K).

To identify immune populations associated with improved viral control, we performed flow cytometry at 8-DPI (fig. S2). CD8^+^ effector T cells play a central role in the clearance of IAV from the lungs (*26–27*). Although total pulmonary CD8^+^ T cell frequencies and counts were comparable between groups, postpartum dams exhibited a significantly greater proportion of antigen-specific CD8^+^ T cells, with a corresponding trend toward increased cell counts (fig. S3A, Fig. 2A). We next determined the effector capacity of pulmonary antigen-specific CD8^+^ T cells by assessing IFN-γ production. At baseline, we observed no significant differences in the frequency or counts of IFN-γ^+^ antigen-specific CD8^+^ T cells between groups (Fig. 2C). However, following PMA/ionomycin stimulation, antigen-specific CD8^+^ T cells from postpartum lungs exhibited significantly more IFN-γ production than those from nulliparous females (Fig. 2D). Thus, although IFN-γ production was comparable at baseline, antigen-specific CD8^+^ T cells from PR8-infected postpartum lungs exhibited an enhanced capacity for effector cytokine production upon stimulation.

Beyond the enhanced antigen-specific CD8^+^ T cell response, we also observed an increased frequency of γδ^+^ T cells in the lungs of postpartum mice at 8-DPI (Fig. 2E). Consistent with the established capacity of γδ^+^ T cells to produce IL-17A, postpartum dams exhibited increased frequencies and counts of IL-17A^+^ γδ^+^ T cells at baseline (Fig. 2F). These findings suggest that pulmonary γδ^+^ T cells in postpartum dams are poised toward IL-17A production during IAV infection, without requiring additional ex vivo stimulation. This difference was amplified following stimulation with recombinant IL-1β and IL-23 (Fig. 2G). In contrast, we observed no difference in the proportion and counts of CD4^+^ T cells, and no difference in IL-17A-producing CD4^+^ T cells, between nulliparous and postpartum dams (fig. S3, B-D, Fig. 2H). Moreover, following stimulation, the count of IL-17A^+^ CD4^+^ T cells was lower than that of IL-17A^+^ γδ^+^ T cells in both groups, suggesting that γδ^+^ T cells, rather than CD4^+^ T cells, represent the predominant T cell source of pulmonary IL-17A during IAV infection (Fig. 2G-H). Consistent with these findings, we also observed a trend toward increased IL-17A cytokine levels in postpartum lungs at 8-DPI (Fig. 2I). Together, these data identify an enhanced IL-17A-producing γδ^+^ T cell response in postpartum lungs during IAV infection.

To directly test the functional contribution of IL-17A to postpartum protection, we administered an IL-17A-neutralizing or isotype control antibody to nulliparous and postpartum dams prior to, and throughout the course, of lethal PR8 infection (Fig. 2J). Consistent with our flow cytometry findings, IL-17A neutralization markedly reduced survival and increased morbidity in postpartum dams, shifting their disease course toward that observed in IAV-infected nulliparous females (Fig. 2K, fig. S3, E-G). Together, these findings reveal an enhanced antigen-specific CD8^+^ T cell response associated with improved viral clearance and establish IL-17A, largely produced by γδ^+^ T cells, as a critical mediator of postpartum resilience to severe IAV disease.

### The transition into lactation promotes maternal resilience to viral infection

We next sought to determine whether lactation contributes to postpartum protection. To address this, we developed a model to compare postpartum dams in the presence or absence of pups, allowing us to distinguish the contribution of active lactation to postpartum protection. Litters were removed from a subset of postpartum dams on the day of birth, followed by infection with a lethal dose of PR8 at 1 day postpartum (Fig. 3A). Strikingly, postpartum dams whose litters were removed exhibited significantly reduced survival and increased morbidity (Fig. 3B, fig. S4, A-C). By 8-DPI, these dams also exhibited significantly higher pulmonary viral burdens, indicating impaired viral clearance (Fig. 3C) and reduced pulmonary IL-17A levels (Fig. 3D). Together, pup removal diminished the enhanced disease resistance observed in postpartum dams, shifting disease severity, viral clearance and pulmonary IL-17A toward levels observed in nulliparous females. These findings suggest that pup-dependent features of lactation are required to maintain maternal resilience to viral infection.

**Fig. 3.**
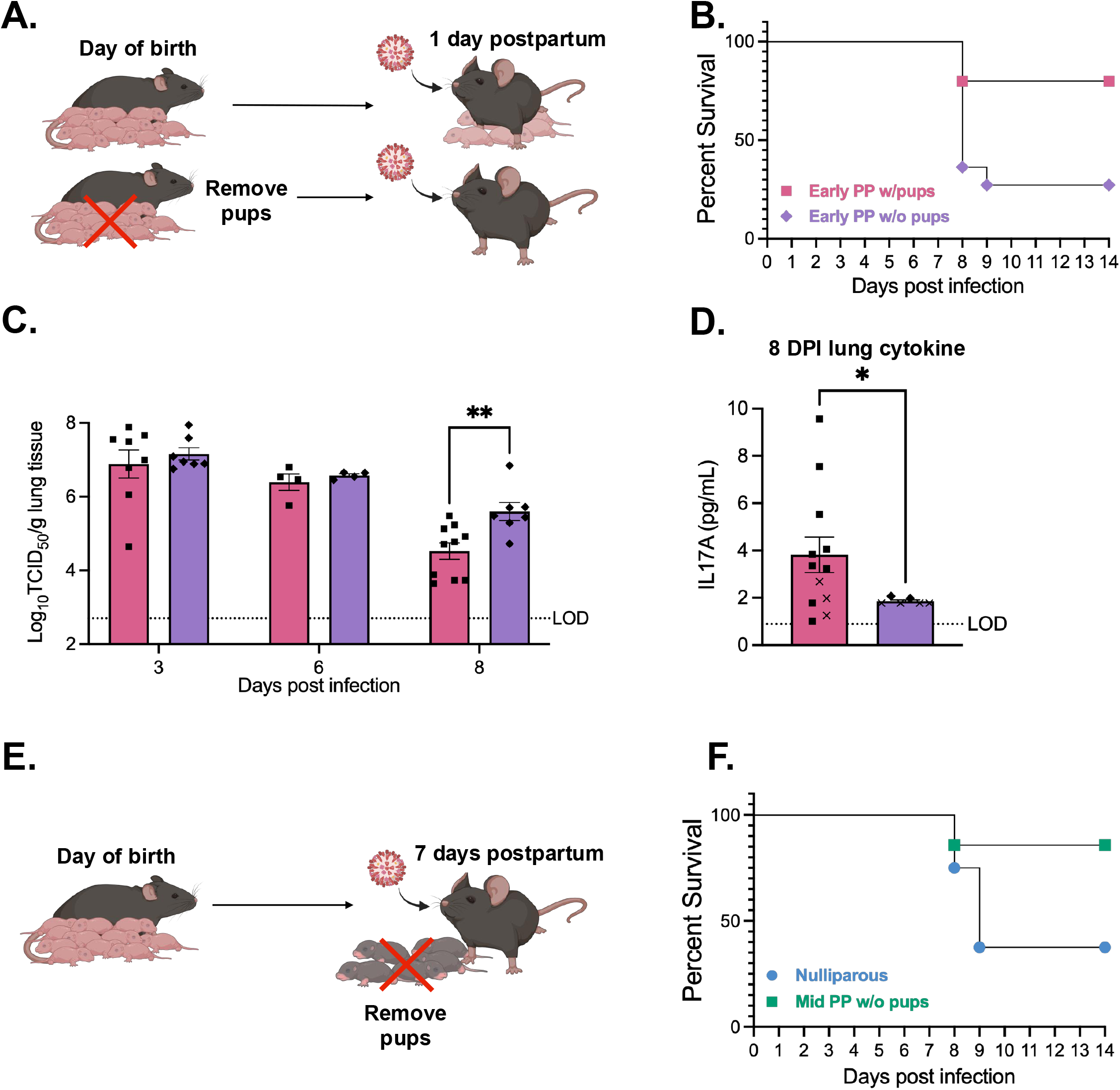
The transition into lactation promotes maternal resilience to viral infection. **(A)** Schematic depiction of the experimental model used to assess postpartum resilience independent of litters. On the day of birth, litters were removed from a subset of dams followed by intranasal inoculation with a lethal dose of PR8 virus at 1-day postpartum and **(B)** daily monitoring for changes in survival (n = 10-11/group) **(C)** Postpartum dams with and without pups were euthanized at 3-, 6-, and 8-DPI to measure viral titers in right lung tissue homogenates via TCID_50_ assay (n = 4-10/DPI and group). **(D)** Left lung homogenates were analyzed for the concentration of IL-17A via LEGENDplex assay (n = 6-12/group). **(E)** Schematic depiction of the experimental model used. Dams gave birth and lactated litters for 7 days. At 7 days postpartum (mid PP), litters were removed from lactating dams followed by intranasal inoculation with a lethal dose of PR8 virus. **(F)** Survival curve of mid PP dams without pups and nulliparous female controls (n = 7-8/group). Dashed lines indicate the limit of detection (LOD) for each assay. Error bars represent the mean ± SEM from 2-4 independent experiments with individual mice indicated by symbols and “x’s” representing moribund mice. *\*P* < 0.05, ^*\*\**^*P* < 0.01. Data were analyzed using two-way ANOVA with Tukey’s multiple comparisons test (C), Unpaired Welch’s t test (D), and Mantel-Cox test for survival analysis, P value = 0.0184 (B), P value = 0.0897 (F).

We next asked whether transition into the lactational state, rather than continued pup presence, was sufficient for postpartum protection. To do this, litters were removed after 7 days of lactation (mid-postpartum), immediately before lethal PR8 infection, with age-matched nulliparous females used as controls (Fig. 3E). Strikingly, mid-postpartum dams that had undergone a period of active lactation exhibited improved survival compared with nulliparous females despite the absence of pups throughout infection (Fig. 3F). Active lactation is accompanied by increased maternal food intake and milk production, which contribute to body mass, and rapidly change following pup removal (*28–29*). Accordingly, mid-postpartum dams without pups exhibited similar changes in body mass to nulliparous females by 8-DPI (fig. S4D). Despite this, mid-postpartum dams without pups better maintained their core body temperature and exhibited fewer clinical signs of disease at 5- and 8-DPI (fig. S4, E-F). Together, our findings suggest that transition into the lactational state is sufficient to establish features of postpartum resistance that are maintained even after the removal of pups.

### Postpartum protection against viral infection emerges through integration of multiple lactation-associated signals

To identify specific factors associated with lactation that contribute to maternal protection, we first determined whether pup-directed maternal caregiving alone was sufficient to confer protection against severe IAV disease. To address this, pregnant dams at embryonic day 15.5 were cohoused with age-matched nulliparous females (“aunties”), allowing aunties to interact with pups beginning at birth. Thereafter, aunties and dams at 1 day postpartum were infected with a lethal dose of PR8 (Fig. 4A). To confirm the establishment of pup-directed caregiving behavior, we performed pup retrieval assays at 0 DPI, prior to infection (1 day postnatal) and at DPI, during the onset of disease (6 days postnatal). Aunties retrieved pups significantly faster than nulliparous controls without prior pup exposure, demonstrating the establishment of maternal caregiving behavior (Fig. 4B). Notably, aunties maintained pup retrieval behavior at 5-DPI, demonstrating that caregiving persisted during IAV disease. Despite the maternal caregiving behavior, aunties exhibited substantially reduced survival and increased morbidity following PR8 infection compared with cohoused postpartum dams (Fig. 4C, fig. S5, A-C). Together, these findings demonstrate that pup-directed maternal caregiving alone is insufficient to recapitulate postpartum resilience to lethal IAV infection and point instead to other physiological features associated with the postpartum lactational state as important determinants of maternal defense.

**Fig. 4.**
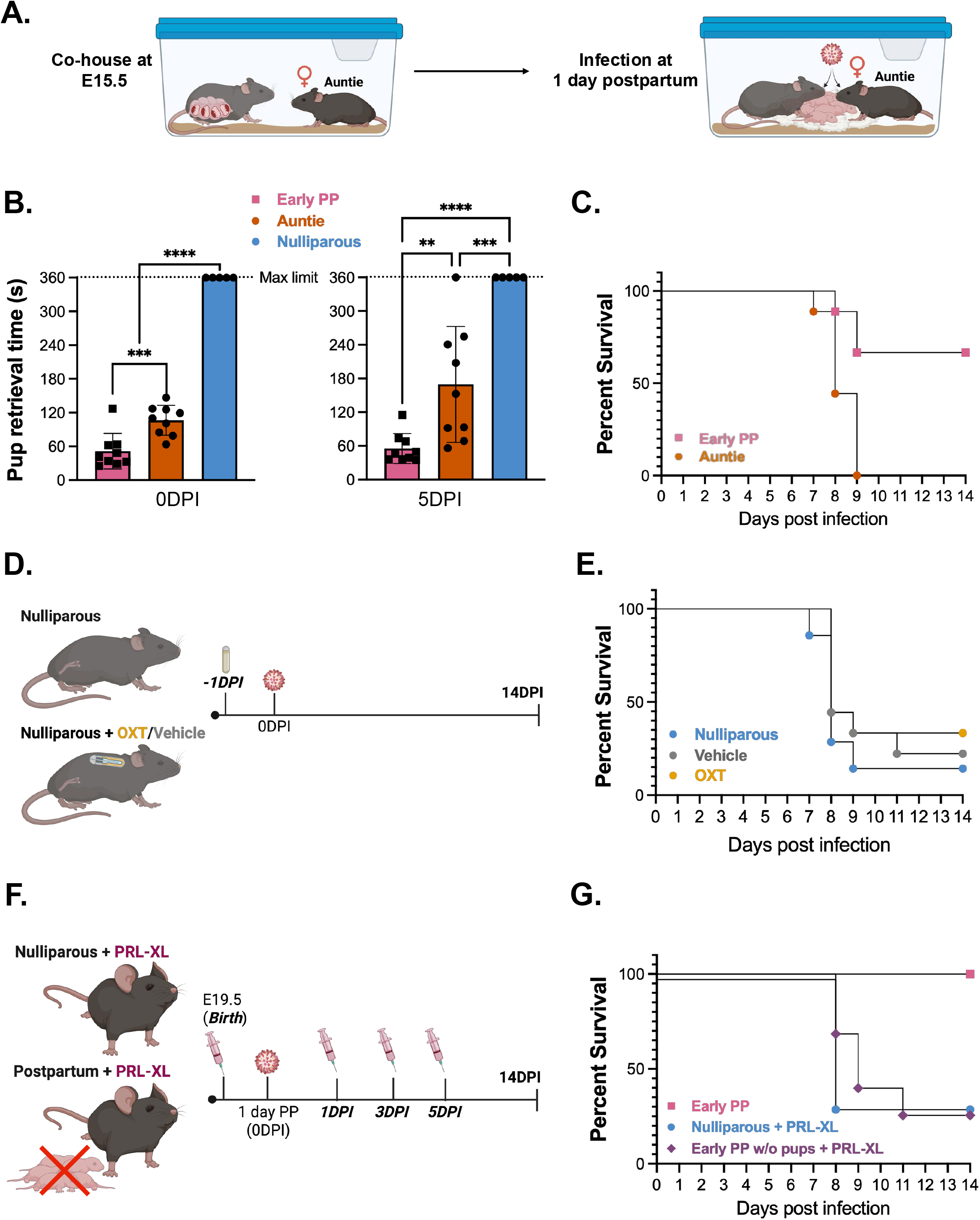
Postpartum protection against viral infection emerges through integration of multiple lactation-associated signals. **(A)** Schematic of the experimental model used to assess caregiving behavior and viral disease outcomes in nulliparous aunties. At embryonic day 15.5, pregnant dams and nulliparous aunties were co-housed, followed by birth and subsequent intranasal inoculation with a lethal dose of PR8 virus at 1-day postpartum in both mice. **(B)** Pup retrieval times for the assessment of maternal behavior are shown in postpartum dams, aunties, and nulliparous controls at 0-DPI (1-day postpartum) and 5-DPI (6-days postpartum). Dashed lines indicate the maximum time limit of the pup retrieval assay (6 minutes, 360 seconds) (n = 5-9/group). **(C)** Survival curve of postpartum dams and aunties (n = 9/group). **(D)** Schematic depiction of the experimental model used for oxytocin (OXT) administration and infection. Nulliparous females received subcutaneous implantation of osmotic pumps for continuous OXT or sterile saline (vehicle) release, with nulliparous females that did not receive osmotic pump implantation used as controls. Thereafter, both groups of females were intranasally inoculated with a lethal dose of PR8 virus and **(E)** survival outcomes were monitored daily (n = 7-9/group). **(F)** Schematic depiction of the experimental model used for exogenous prolactin-XL treatment. On the day of birth, litters were removed and postpartum dams and nulliparous females were treated with prolactin-XL (PRL-XL) via i.p. injections. Thereafter, both groups were intranasally inoculated with a lethal dose of PR8 virus and received subsequent i.p. injections of PRL-XL through 5-DPI. **(G)** Survival curve of postpartum dams with pups, nulliparous females and postpartum dams without pups given PRL-XL (n = 2-7/group). Error bars represent the mean ± SEM from 3 independent experiments with individual mice indicated by symbols. *\*P* < 0.05, ^*\*\**^*P* < 0.01, ^*\*\*\**^*P* < 0.001, ^*\*\*\*\**^*P* < 0.0001. Data were analyzed using ordinary one-way ANOVA with Tukey’s test (B), and Mantel-Cox test for survival analysis, P value = 0.0027 (C), P value = 0.5190 (E), P value = 0.2364 (G).

In addition to postnatal caregiving, the lactational period is accompanied by substantial hormonal changes, including increased oxytocin and prolactin signaling that support milk let-down and production, respectively (*30*). Importantly, both hormones have documented immunomodulatory functions, raising the possibility that lactation-associated hormonal signaling contributes to postpartum resilience during viral infection (*31–33*). We first asked whether oxytocin alone was sufficient to recapitulate the enhanced resistance to severe IAV disease observed in postpartum dams. To test this, nulliparous females received continuous oxytocin or sterile saline (vehicle) through subcutaneously implanted osmotic pumps, allowing sustained oxytocin delivery given its short circulating half-life. 24 hours after pump implantation, females were infected with a lethal dose of PR8, with untreated nulliparous females without osmotic pumps included as additional controls (Fig. 4D). Despite sustained oxytocin administration, oxytocin-treated nulliparous females exhibited comparable survival and morbidity to vehicle-treated and untreated nulliparous controls (Fig. 4E, fig. S5, D-F). These findings demonstrate that oxytocin signaling alone is insufficient to recapitulate postpartum resilience to lethal IAV infection.

Given the central role of prolactin in lactation, we next tested whether prolactin signaling was sufficient to promote maternal resilience to viral infection. To test this, we administered an engineered long-lasting prolactin (Prolactin-XL) (*34*) to nulliparous females one day prior to infection with a lethal dose of PR8, followed by subsequent hormone administration at 1, 3, and 5-DPI (Fig. 4F). Prolactin-treated nulliparous females remained susceptible to lethal infection, with significantly reduced survival compared with postpartum lactating dams (Fig. 4G). Thus, prolactin signaling alone was insufficient to recapitulate the enhanced resilience observed during the postpartum period. We next asked whether prolactin could restore postpartum protection following pup removal. Litters were removed from dams on the day of birth, followed by administration of Prolactin-XL. At 1 day postpartum, dams were infected with a lethal dose of PR8 and received subsequent prolactin administration at 1, 3, and 5-DPI (Fig. 4F). Despite restoration of prolactin signaling, and visible nipple engorgement (data not shown), postpartum dams without litters exhibited survival comparable to prolactin-treated nulliparous females and substantially reduced survival compared with postpartum lactating dams (Fig. 4G). Both prolactin-treated groups exhibited greater loss of body mass than postpartum dams with litters (fig S5G). Notably, at peak disease, prolactin-treated postpartum dams without litters exhibited less severe hypothermia and fewer clinical signs of disease than prolactin-treated nulliparous females, although these measures remained more pronounced than in postpartum dams with litters (fig. S5, H-I). These findings suggest that prolactin may modulate specific features of disease severity within the postpartum context but is insufficient on its own to restore the full protective phenotype.

Together, our studies demonstrate that postpartum protection from viral infection cannot be attributed to any single feature of the lactational period examined here. Rather, our findings support a model in which maternal resilience likely emerges through the integration of multiple physiological signals associated with the transition into the active postpartum lactational state.

### Postpartum resilience extends to highly pathogenic avian influenza H5N1 infection

Highly pathogenic avian influenza (HPAI) H5N1 virus is a growing public health concern due to ongoing cross-species transmission and its potential to cause severe disease in humans (*35–36*). Given the increasing clinical relevance of H5N1 and the enhanced resilience observed following PR8 infection, we sought to determine whether postpartum protection extends to HPAI H5N1. Lactating dams were intranasally inoculated with HPAI H5N1 at 1 day postpartum, with age-matched nulliparous females used as controls. As previously described, litters were culled to six pups on postnatal day 3 to standardize caregiving load. Remarkably, postpartum dams exhibited improved survival following H5N1 infection, with 54% surviving compared with only 20% of nulliparous females (Fig. 5A). Consistent with this survival advantage, postpartum dams better maintained their body mass and core body temperature throughout the course of infection, including during peak disease at 8-DPI (Fig. 5B-C). Postpartum dams also exhibited fewer clinical signs of HPAI H5N1 disease compared with nulliparous females, further demonstrating robust postpartum resilience to a highly pathogenic influenza virus (Fig. 5D). Lastly, we assessed maternal behavior and found that H5N1-infected postpartum dams maintained pup retrieval behavior, with retrieval times below the maximum assay limit and comparable to those observed in PR8-infected postpartum dams (Fig. 5E). Moreover, pup retrieval times did not significantly differ between 5- and 8-DPI, demonstrating that postpartum dams maintained maternal caregiving even as H5N1 disease progressed to peak severity. Together, these findings demonstrate that postpartum resilience extends beyond PR8 infection to a highly pathogenic influenza virus while preserving maternal caregiving during severe disease.

**Fig. 5.**
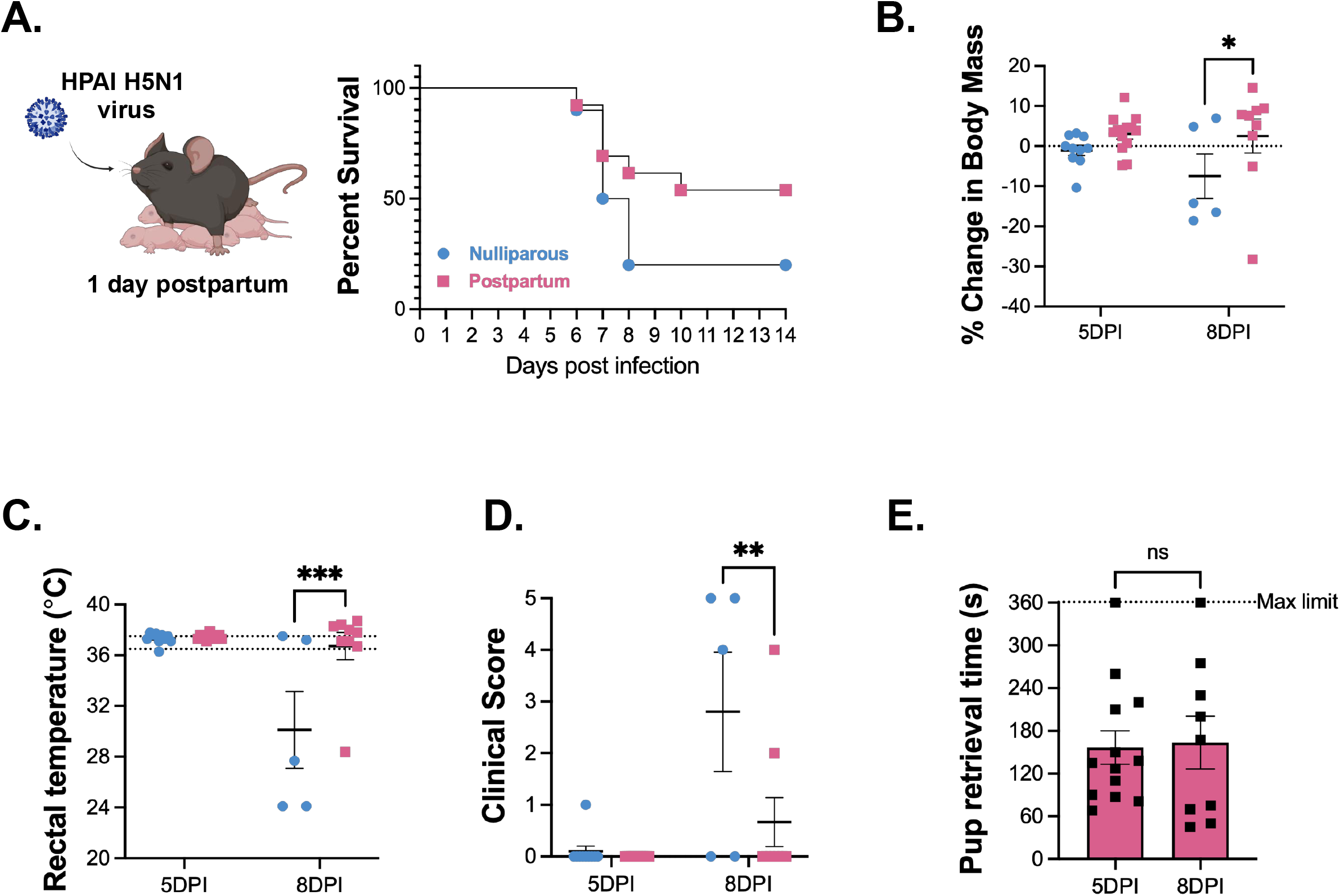
Postpartum resilience extends to highly pathogenic avian influenza (HPAI) H5N1 virus. **(A)** At 1-day postpartum dams were intranasally inoculated with 10 TCID_50_ of high pathogenicity avian influenza (HPAI) H5N1 with age-matched nulliparous females used as controls. **(A)** Mice were monitored daily for changes in survival, and **(B)** starting body mass, **(C)** rectal temperature, and **(D)** clinical signs of disease were analyzed at 5-(onset of disease) and 8-DPI (peak disease) (n = 10-13/group). For percent change in body mass, dashed lines at 0 indicate starting body mass. For rectal temperature, dashed lines indicate standard core body temperature of laboratory mouse (36.5 to 37.5 °C). **(E)** Pup retrieval times are shown for H5N1-infected postpartum dams at 5- and 8-DPI. The maximum limit of the pup retrieval assay (6 minutes, 360 seconds) is indicated by a dashed line (n = 9-13/group). Error bars represent the mean ± SEM from 2 independent experiments with individual mice indicated by symbols. *\*P* < 0.05, ^*\*\**^*P* < 0.01, ^*\*\*\**^*P* < 0.001, ns *P* > 0.05. Data were analyzed using two-way ANOVA with Fisher’s LSD (B-D), paired t tests (E), and Mantel-Cox for survival analysis, P value = 0.1161 (A).

## DISCUSSION

Maternal resilience to physiological and infectious stressors is essential for sustaining both maternal and newborn survival. Given the substantial physiological and caregiving demands of motherhood, the postpartum period has likely been shaped by strong evolutionary pressures to preserve maternal health while supporting offspring survival. Our study provides an extensive exploration of the murine postpartum and lactational periods that supports this concept. Our findings challenge the idea that maternal physiology simply returns to a nonpregnant baseline following childbirth and instead establish the postpartum lactational period as a distinct physiological state adapted to combat severe viral infection, consistent with clinical and epidemiological observations (*37–39*). Notably, infected postpartum dams maintained maternal caregiving behavior while combating severe viral disease, demonstrating that enhanced resistance to infection does not come at the expense of offspring care. Together, these findings suggest that postpartum physiology is adapted to balance the simultaneous demands of offspring care and resistance to severe viral disease.

We demonstrate distinct maternal immune-mediated mechanisms of antiviral protection in the lung following lethal viral infection and link specific cytokine signaling to improved viral clearance and disease outcomes in postpartum dams. Interestingly, we observed increased pulmonary γδ T cells in postpartum mice late in infection. Although γδ T cells are enriched at mucosal sites (*40*) and are often characterized as rapid responders during infection (*41*), their increased abundance at this later stage of IAV infection suggests that the postpartum state may alter the magnitude or kinetics of the pulmonary γδ T cell response. These postpartum-associated changes in innate and innate-like immunity may shape subsequent adaptive responses, potentially contributing to the enhanced antigen-specific T cell responses observed during infection. Together, these findings reveal distinct, but potentially coordinated, changes across innate and adaptive immunity that contribute to enhanced protection against lethal IAV infection during the postpartum period.

In humans, sustained lactation is associated with reduced risks of stroke, metabolic disease, and cardiovascular disorders (*42–45*). Our findings suggest that enhanced antiviral resilience may represent an additional, previously unrecognized benefit associated with the lactational state. Our findings identify the transition into lactation as a critical window for establishing maternal antiviral resilience and suggest that protection emerges through the integration of multiple lactation-associated physiological signals. Importantly, establishment of active lactation was sufficient to induce postpartum-derived protection even in the absence of continued pup suckling during infection. Conversely, no individual feature of the lactational period examined here was sufficient to recapitulate maternal resilience to viral infection. Rather, postpartum resilience may emerge through the integration of multiple physiological signals associated with the transition into the lactational state.

Finally, we demonstrate that postpartum resilience extends to another clinically relevant virus, highly pathogenic avian influenza H5N1. Beyond establishing the breadth of postpartum resilience across distinct influenza virus infections, these findings demonstrate that enhanced protection is maintained even in the context of a highly pathogenic viral challenge.

Collectively, our study establishes the postpartum period as an important and understudied frontier in fourth-trimester infectious disease. More broadly, this work provides a framework for understanding how maternal immunity remains adapted to the demands of reproduction even after birth, supporting host defense during a period in which maternal health is essential for both maternal and offspring survival.

## Acknowledgments

The data for this manuscript were generated in the Penn Cytomics and Cell Sorting Shared Resource Laboratory at the University of Pennsylvania (RRID:SCR_022376). Penn Cytomics is partially supported by the Abramson Cancer Center NCI Grant (P30 016520). We thank the NIH Tetramer Core Facility (contract number 75N93020D00005) for providing influenza tetramers. We would like to thank Daniel Yong, as well as the Mike Hogan, Laurence Eisenlohr, and Zoltan Arany laboratories for discussions about these data. Schematic representations in this manuscript were created using Biorender.com.

## Author contributions

Conceptualization: ADC, KAJ Methodology: ADC, TME, MCA, KAJ Investigation: ADC, TME, MCA, EAK, PH, CS, NE, MM, KK, SH Visualization: ADC, MCA Funding acquisition: KAJ Project administration: ADC, KAJ Supervision: PAS, KAJ Writing – original draft: ADC, KAJ Writing – review & editing: ADC, TME, MCA, EAK, PH, CS, NE, MA, KK, PAS, SH, KAJ

## Diversity, equity, ethics, and inclusion [optional]

In the Jurado laboratory, we center equity and inclusion. We recognize, appreciate and celebrate the distinct backgrounds and diverse perspectives that drive our science forward. We, as a team, nurture a collaborative environment that fosters integrity and inclusion in tandem with our rigorous research pursuits. We foster a team culture where everyone feels safe, respected, and empowered.

## Data, code, and materials availability

All data needed to evaluate the conclusions of this paper are provided in the manuscript and supplementary materials. Raw data are available from the corresponding author (KAJ) upon request.

## Supplementary Materials

Figs. S1 to S#5

## Materials and Methods

### Mice

Wild-type (WT) C57BL/6 (B6) mice were procured from The Jackson Laboratory. For timed mating pregnancies, 7-to 12-week-old females were co-housed with male breeder mice. Age-matched females that were co-housed with male mice but did not show a copulation plug or did not get pregnant were used as nulliparous controls. WT B6 nulliparous females with the same birth date and from the same delivery as pregnant/postpartum dams were used as Aunties. All postpartum dams with pups present in the cage had litters culled down to six pups on postnatal day 3 to standardize caregiving load. All mice were maintained at the University of Pennsylvania under specific pathogen free conditions, following the University of Pennsylvania approved IACUC protocol (#806749).

### Pup retrieval assay for maternal behavior

Postpartum dams were placed in a clean cage to acclimate for 10 minutes, while pups were removed out from nest onto a heating pad. After 10 minutes, 5 pups per dam were returned to the home cage and randomly distributed away from the nest, with any remaining pups on the heating pad. Postpartum dams were placed back into the home cage and the time it takes for dams to retrieve all 5 pups and place them back into the nest was recorded. Pup retrieval was defined as picking up the pup in the mouth and returning it to the nest (*23–24*). Nulliparous females with no prior exposure to pups at all do not exhibit a pup retrieval response.

### H1N1 virus

Influenza A virus H1N1 strain A/Puerto Rico/8/1934 (influenza A) was obtained from the Hensley Laboratory at the University of Pennsylvania.

### H1N1 infection

Influenza A PR8 virus was diluted to 100 TCID_50_ (sub-lethal) and 250 TCID_50_ (lethal) in 1X DPBS for all infections. Mice were anesthetized using isoflurane inhalation and were given 50µL of inoculum intranasally (25µL per nostril). Mice were monitored out to 14 days post-infection, with body mass, rectal (core) temperature, and clinical score recorded daily. Clinical signs were defined as piloerection, hunched posture, dyspnea, absence of an escape response, and termination due to 30% reduction in body mass or temperature below 30°C. Mice that displayed 4 clinical signs or more were humanely euthanized.

### H5N1 virus

The virus was propagated from our stock isolate virus (A/dairy cattle/Texas/24-008749-002/2024 HPAI H5N1 B3.13 high pathogenicity) in MDCK (Madin-Darby Canine Kidney) cells. The MDCK cells were grown in Dulbecco’s modified Eagle’s medium DMEM, supplemented with 35% BSA (bovine albumin solution), 5% Sodium Bicarbonate, and 5mL of Penicillin/Streptomycin. The virus was sequenced after propagation and found to match the input strain.

### H5N1 infection

Mice were housed in the Animal Biosafety Level 3 Facility at the University of Pennsylvania under the standard dark/light cycle, ambient temperature, humidity, and within the Allentown individual ventilated BCU2 caging system adhering to the University of Pennsylvania’s approved ABSL-3 IBC and IACUC protocols (#807017). Mice were anesthetized with ketamine-xylazine and infected with 10 TCID_50_ of high pathogenicity H5N1 intranasally diluted within 40uL PBS. Mice were monitored daily and scored for clinical disease, as well as body weight and temperature being measured. At clinical or experimental endpoint (Day 14 post infection), mice were euthanized by isoflurane.

### TCID_50_ assay

To assess influenza viral burden, right lung lobes were harvested from infected mice at 3-, 6-, and 8-days post-infection. Lungs were harvested into serum-free 1X MEM, homogenized, and aliquoted into 1.5mL Eppendorf tubes. MDCK cells were seeded at 50,000 cells/well in 96-well flat bottom plates. MDCK cells were maintained in MEM supplemented with 10% heat-inactivated FBS. At 24 hours post-seeding, plates were washed twice with 200µL warm MEM and 180µL of TCID_50_ medium (containing 50mL serum-free 1X MEM, 50µL gentamycin sulfate, 50µL TPCK-trypsin, 250µL HEPES) was added to each well. Homogenized samples were briefly vortexed and spun and 20µL of each sample was added to the first column of the 96-well plate with infection medium in quadruplicate. 10-fold dilutions were performed across the plate, with the last well remaining as a control. Plates were incubated at 37°C for 3 days.

Individual wells were visually evaluated for cytopathic effect (CPE) and the Reed and Muench method was used to calculate TCID_50_/mL titer for each sample.

### Single-cell isolation from lung tissue

Lungs were harvested, minced, and digested with 2 mg/mL of Collagenase Type IV (Gibco, Catalog No. 17104019) and 1 mg/mL of DNAse I (Millipore Sigma, Catalog No. 10104159001) in Complete Media [RPMI 1640 (Invitrogen), 1% of 100X GlutaMAX (Invitrogen), 1% of 100X NEAA (Gibco), 20 mM HEPES (Invitrogen), 0.1% of 100X BME (Millipore Sigma), 1% of 100X Pen/Step (Corning)] for 20 minutes at 5% CO^2^, 37°C. The enzymes were neutralized by adding Complete Media containing 10% FBS. Lung digests were strained through a 70-μm nylon mesh and red blood cells were lysed with Ammonium-Chloride-Potassium (ACK) Lysing Buffer (Quality Biological Inc.). Lung samples were resuspended in 30% Stock Isotonic Percoll [90 mL Percoll (Cytiva; Catalog No. 17089101) and 10 mL 10X PBS (Invitrogen)] diluted with HBSS (Invitrogen). Cells in 30% Stock Isotonic Percoll were centrifuged at 1200 xg for 20 min at 20°C. Lung-isolated cells were resuspended, cell counts were determined with a hemacytometer, and 1 × 10^6^ cells were used for cytokine stimulation followed by flow cytometry staining.

### Cytokine stimulation

For the characterization of IL-17A and IFNγ synthesis by immune cells in the lungs, 1 × 10^6^ lung-isolated cells were plated in a flat-bottom 96-well plate in 200 μL of stimulation media [RPMI 1640 (Invitrogen), 10% FBS (heat-inactivated), 1% of 100X GlutaMAX (Invitrogen), 1% of 100X NEAA (Gibco), 20 mM HEPES (Invitrogen), 0.1% of 100X BME (Millipore Sigma), 1% of 100X Pen/Step (Corning)], stimulated with rIL-23 (10 ng/mL) (BioLegend, Catalog No. 589002) and rIL-1B (10 ng/mL) (BioLegend, Catalog No. 575102) or PMA/Ionomycin (Invitrogen, Catalog No. 00-4970-03), and incubated with GolgiStop (1:1000) (BD Biosciences, Catalog No. 554724) for 4 hours at 5% CO^2^, 37°C.

### Flow cytometry staining

Cells were initially stained with Live/Dead Fixable Aqua Dead Cell Stain (Invitrogen, Catalog No. L34966) diluted 1:1000 in PBS for 20 minutes at 4°C. To identify lymphoid immune cells, cells were surface-stained (30 minutes; 4°C) with Pacific Blue-conjugated anti-CD3e (BioLegend; Clone 17A2), Brilliant Violet 605-conjugated anti-CD4 (BioLegend; Clone GK1.5), FITC-conjugated anti-CD8a (BioLegend; Clone 53-6.7), PerCP-eFlour 710-conjugated anti-γδTCR (Invitrogen; Clone GL3), PE-conjugated anti-CD45 (BioLegend; Clone 30-F11), APC-Cyanine7-conjugated anti-CD25 (BioLegend; Clone PC61), and BUV395-conjugated anti-CD19 (BD Biosciences; Clone ID3) in Cell Stain Buffer (BioLegend; Catalog No. 420201) containing 1:100 TruStain FcX™ PLUS (anti-mouse CD16/32) Antibody (BioLegend; Catalog No. 156604). To identify antigen-specific CD8 T cells, cells were surface-stained (30 minutes; 4°C) with Influenza A NP366-374 PE-labeled Tetramer (NIH Tetramer Core Facility), Brilliant Violet 605-conjugated anti-CD3e (BioLegend; Clone 17A2), AF488-conjugated anti-CD4 (BioLegend; Clone GK1.5), APC-conjugated anti-CD8a (BioLegend; Clone 53-6.7), APC-F750-conjugated anti-CD45 (BioLegend; Clone 30-F11), BUV395-conjugated anti-CD19 (BD Biosciences; Clone ID3) in Cell Stain Buffer (BioLegend; Catalog No. 420201) containing 1:100 TruStain FcX™ PLUS (anti-mouse CD16/32) Antibody (BioLegend; Catalog No. 156604). For intracellular staining, cells were fixed and permeabilized using the eBioscience™ Foxp3/Transcription Factor Staining Buffer Set (ThermoFisher Scientific, Catalog No. 00-5523-00) following manufacturer instructions. Cells were intracellularly stained with BV785-conjugated anti-IL-17A (BioLegend; Clone TC11-18H10.1) or BV421-conjugated IFNγ (BioLegend; Clone XMG1.2). Flow cytometry was performed on a LSR Fortessa B (BD Biosciences). Analysis was performed using FlowJo software version 10.10.1.

### LEGENDplex cytokine assay

Lungs were harvested and homogenized in Homogenizing Media [1 tablet of cOmplete, Mini Protease Inhibitor Cocktail (Roche; Catalog No. 11836153001) to 15 mL of DPBS (Corning; Catalog No. 21-031-CV)]. Homogenate was centrifuged and supernatant was collected and frozen at −80°C. Cytokine concentrations were determined by following manufacturer instructions using a custom LEGENDplex (BioLegend) bead-based immunoassay for the following cytokines: IL-17A, TNF-α, IL-12p40, IL-1β, IL-12p70, IL-23, GM-CSF, IL-10, IL-6, IL-27, IFN-γ, IFN-β, IFN-α2, IL-1α. Samples were analyzed on a LSR Fortessa B (BD Biosciences). Analysis was performed using the LEGENDplex (BioLegend) online analysis platform (https://legendplex.qognit.com/).

### Administration of IL-17A blocking antibody

Nulliparous and early postpartum dams were administered anti-IL-17A (~213µg per mouse; #BE0173, clone 17F3, BioXCell) or anti-IgG1κ (~205µg per mouse; #BE0083, clone MOPC-21, BioXCell) in 200µL of 1X DPBS intraperitoneally the day of birth (E19.5; 1 day prior to influenza PR8 infection) and every other day out to 7-days post-infection as previously described (*46*). Mice were monitored daily out to 14 days post-infection as described previously.

### Murine hormone administration

For oxytocin hormone treatment, nulliparous females were anesthetized using isoflurane inhalation (3-4%) and received surgical operation to implant osmotic mini pumps (Alzet; model 1002, CA) into the subcutaneous tissue. Osmotic mini pumps contained 100µL of sterile physiological water (vehicle control group) or 100µL of oxytocin acetate salt hydrate (100µg total, 6µg/day for 14 days; #O6379, Sigma-Aldrich) (oxytocin group). All implantations of osmotic mini pups took place 1 day prior to influenza PR8 infection.

An engineered long-lasting prolactin (prolactin-XL) generated by the Wyss institute at Harvard University was used for prolactin hormone treatment described below (*34*). Nulliparous and postpartum dams without litters were administered prolactin-XL (5mg/kg) in 100µL of 1X DPBS intraperitoneally the day of birth (E19.5; 1 day prior to influenza PR8 infection) and every other day out to 5-days post-infection. On the day of birth, litters were immediately removed from the dam home cage and euthanized, followed by subsequent hormone administration.

### Statistical analysis

All graphs were plotted using GraphPad Prism software (version 11.0.2) and statistical analyses were performed using GraphPad Prism. Data are expressed as mean with standard error with scatter plots showing individual values. Sample size, number of individual experiments, and statistical tests used for each experiment are detailed in figure legends. Survival analyses were performed using the Mantel-Cox test. Statistical differences in changes in body mass and core rectal temperature were determined with multiple unpaired t tests or two-way ANOVA with Fisher’s LSD, and differences in clinical scores (categorical) were determined by Mann-Whitney tests, with Holm-Šidák correction when appropriate. All viral loads were analyzed using two-way ANOVA with Tukey’s multiple comparisons test. Frequencies and absolute number of immune cells were analyzed using unpaired Welch’s t test or Mann-Whitney tests where appropriate. Pup retrieval for maternal behavior times were analyzed using ordinary one-way ANOVA with Tukey’s test or paired t tests where appropriate. p < 0.05 was considered as significant. ns = non-significant, *\*P < 0*.*05*, ^*\*\**^*P < 0*.*01*, ^*\*\*\**^*P < 0*.*001*, ^*\*\*\*\**^*P < 0*.*0001*.

