## Supplementary Figures for "The transition to postpartum lactation establishes a state of maternal antiviral resilience"

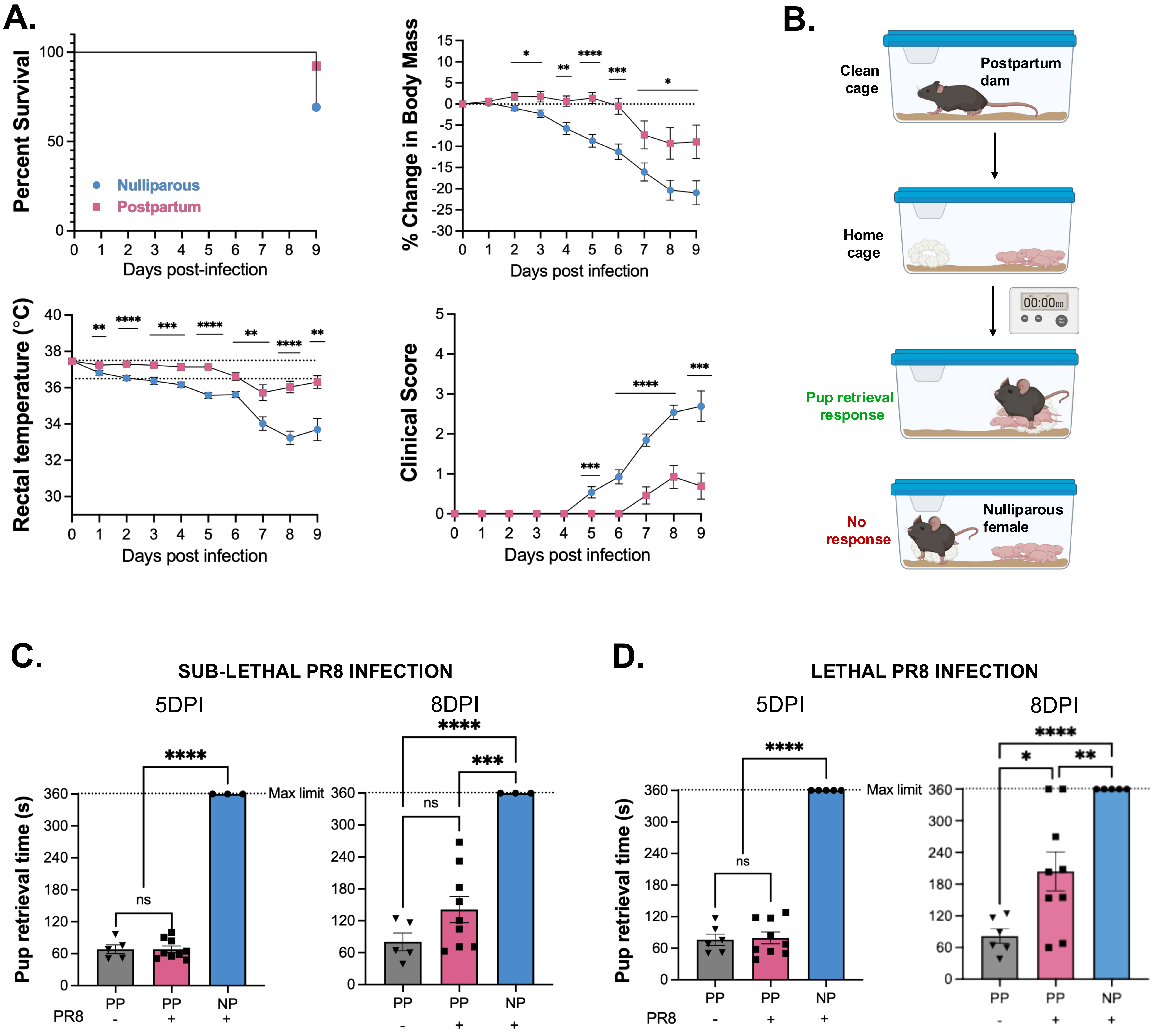
A.
Percent Survival
Nulliparous
Postpartum
Days post-infection
% Change in Body Mass
Days post infection
Rectal temperature (°C)
Days post infection
Clinical Score
Days post infection
B.
Clean cage
Postpartum dam
Home cage
00:00
Pup retrieval response
No response
Nulliparous female
C.
SUB-LETHAL PR8 INFECTION
5DPI
8DPI
Pup retrieval time (s)
Max limit
ns
PP
NP
PR8
D.
LETHAL PR8 INFECTION
5DPI
8DPI
Pup retrieval time (s)
Max limit
ns
PP
NP
PR8

Fig. S1.

**Fig. S1. Postpartum mice display resilience to sub-lethal IAV infection and maintain maternal caregiving behavior.**

**(A)** Postpartum dams were intranasally inoculated with 100 $TCID_{50}$ (sub-lethal dose) of influenza A/PR/8/34 (PR8) at 1-day post-birth. Dams were monitored daily for 14 days to assess changes in survival, starting body mass, rectal temperature, and clinical score (n = 13/group). For percent change in body mass, dashed lines at 0 indicate starting body mass. For rectal temperature, dashed lines indicate standard core body temperature of laboratory mouse (36.5 to 37.5 °C). **(B)** Schematic representation of the pup retrieval assay to assess maternal behavior. **(C and D)** Pup retrieval times for the assessment of maternal behavior are shown at 5- and 8-DPI in nulliparous (NP) females and postpartum (PP) dams with or without sub-lethal (n = 3-9/group) **(C)** and lethal (n = 5-9/group) **(D)** PR8 infection. Dashed lines indicate the maximum time limit of the pup retrieval assay (6 minutes, 360 seconds). Error bars and individual symbols represent the mean ± SEM for 3-4 independent experiments. Individual mice are indicated by symbols (C and D). *$P < 0.05$, **$P < 0.01$, ***$P < 0.001$, ****$P < 0.0001$, ns $P > 0.05$. Data were analyzed using multiple unpaired t tests for change in body mass and rectal temperature and multiple Mann-Whitney tests for clinical score (A). Survival analysis was performed with Mantel-Cox test, P value = 0.1665 (A). Ordinary one-way ANOVA with Tukey's test was used for analysis of pup retrieval times (C and D).

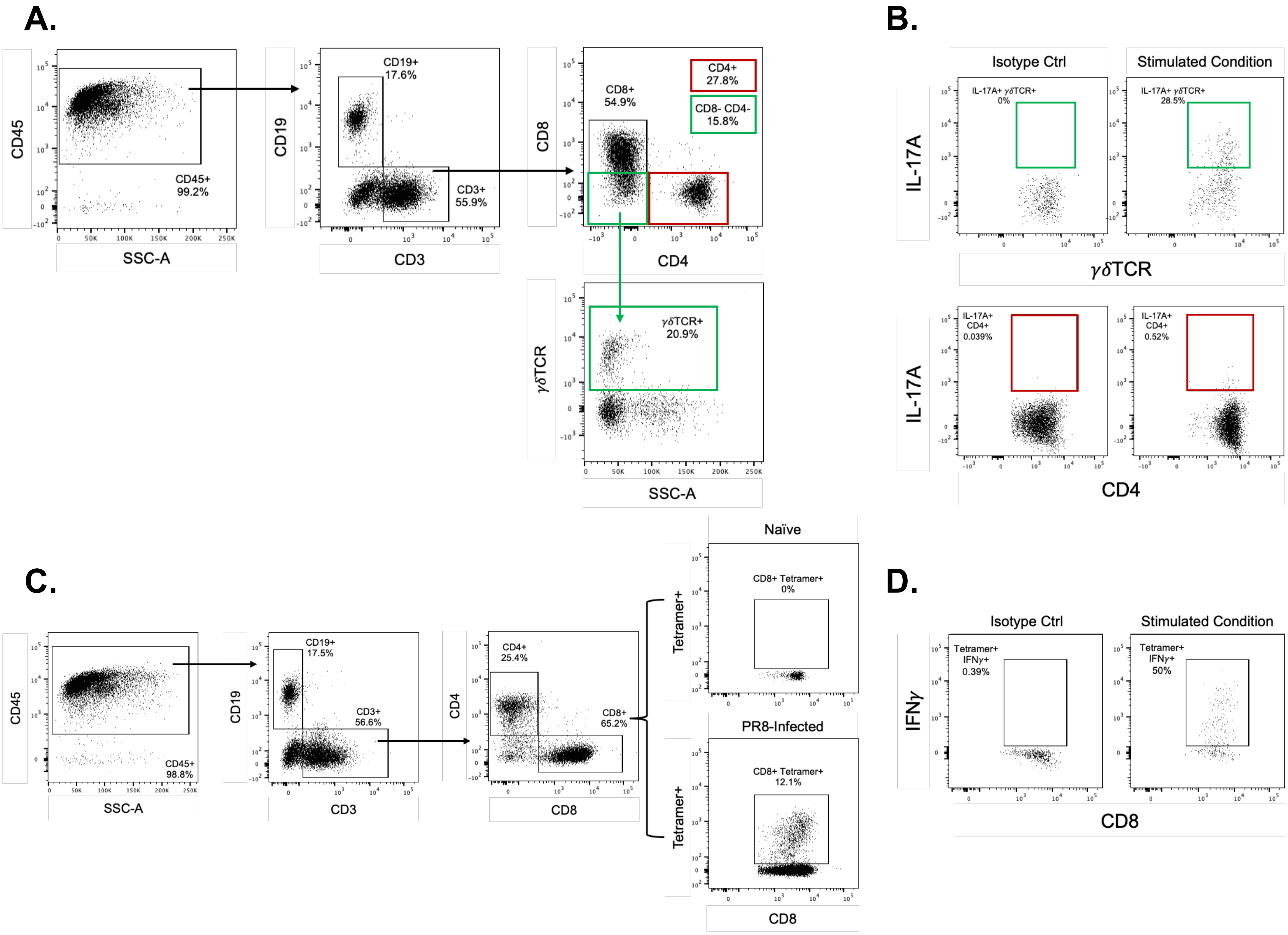

A.
CD45
SSC-A
CD45+
99.2%
CD19
CD3
CD19+
17.6%
CD3+
55.9%
CD8
CD4
CD8+
54.9%
CD4+
27.8%
CD8- CD4-
15.8%
γδTCR
γδTCR+
20.9%
B.
Isotype Ctrl
Stimulated Condition
IL-17A
IL-17A+ γδTCR+
0%
IL-17A+ γδTCR+
28.5%
IL-17A+
CD4+
0.039%
IL-17A+
CD4+
0.52%
C.
CD45+
98.8%
CD19+
17.5%
CD3+
56.6%
CD4+
25.4%
CD8+
65.2%
Naïve
Tetramer+
CD8+ Tetramer+
0%
PR8-Infected
CD8+ Tetramer+
12.1%
D.
IFNγ
Tetramer+
IFNγ+
0.39%
Tetramer+
IFNγ+
50%


**Fig. S2.**

**Fig. S2. Gating strategy for immune cells in the lung.**

Gating strategy for immune cells in the lungs. All populations were gated from single, viable cells. **(A)** γδ T cells (CD45+ CD3+ CD8- CD4- γδ TCR+) and CD4+ T cells (CD45+ CD3+ CD8- CD4+). **(B)** Representative FACS plots of IL-17A cytokine production by γδ T and CD4 T cells after stimulation with recombinant IL-23 and IL-1β and their respective isotype controls. **(C)** Gating strategy to identify antigen-specific CD8 T cells. CD8 T cells (CD45+ CD3+ CD8+). Representative FACS plots of CD8+ $NP_{366\text{-}374}$ tetramer+ T cells from either a naïve or PR8-infected mouse. (D) Representative FACS plot if IFNγ production by CD8 T cells after stimulation with PMA/Ionomycin and respective isotype control.

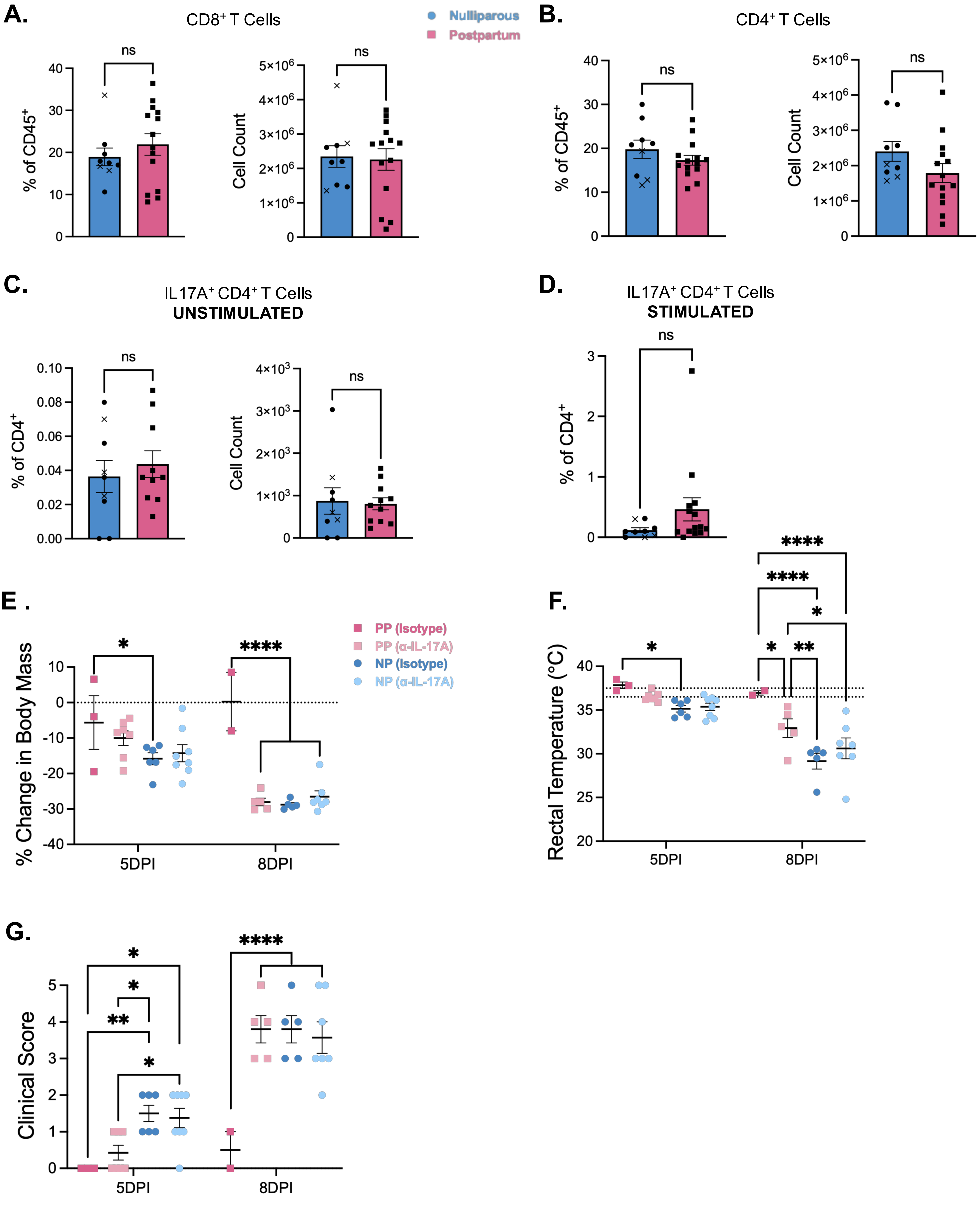
A.
CD8⁺ T Cells
Nulliparous
Postpartum
% of CD45⁺
Cell Count
ns
B.
CD4⁺ T Cells
% of CD45⁺
Cell Count
ns
C.
IL17A⁺ CD4⁺ T Cells
UNSTIMULATED
% of CD4⁺
Cell Count
ns
D.
IL17A⁺ CD4⁺ T Cells
STIMULATED
% of CD4⁺
ns
E .
PP (Isotype)
PP (α-IL-17A)
NP (Isotype)
NP (α-IL-17A)
% Change in Body Mass
5DPI
8DPI
F.
Rectal Temperature (°C)
5DPI
8DPI
G.
Clinical Score
5DPI
8DPI

Fig. S3.

**Fig. S3. Postpartum pulmonary T cells and response to IL-17A depletion.**

Nulliparous and postpartum dams were intranasally inoculated with a lethal dose of PR8 virus. Flow cytometry was performed using live cells from whole lung at 8DPI to assess **(A)** $CD8^+$ and **(B)** $CD4^+$ T cell frequencies and counts (n = 9-14/group). **(C)** Frequency and counts of IL-17A-producing $CD4^+$ T cells are shown during unstimulated conditions and following **(D)** stimulation with IL-1β and IL-23 (n = 9-14/group). **(E)** Change in starting body mass, **(F)** core temperature, and **(G)** clinical signs of disease are shown at 5- and 8-DPI in nulliparous (NP) and postpartum (PP) mice treated with anti-IL-17A or isotype antibody (n = 3-8/group). For percent change in body mass, dashed lines at 0 indicate starting body mass. For rectal temperature, dashed lines indicate standard core body temperature of laboratory mouse (36.5 to 37.5 °C). Error bars represent the mean ± SEM from 3-4 independent experiments with individual mice indicated by symbols and "x's" representing moribund mice. *$P < 0.05$, **$P < 0.01$, ***$P < 0.001$, ****$P < 0.0001$, ns $P > 0.05$. Data were analyzed using Unpaired Welch's t test (A-D) and two-way ANOVA with Fisher's LSD (E-G).

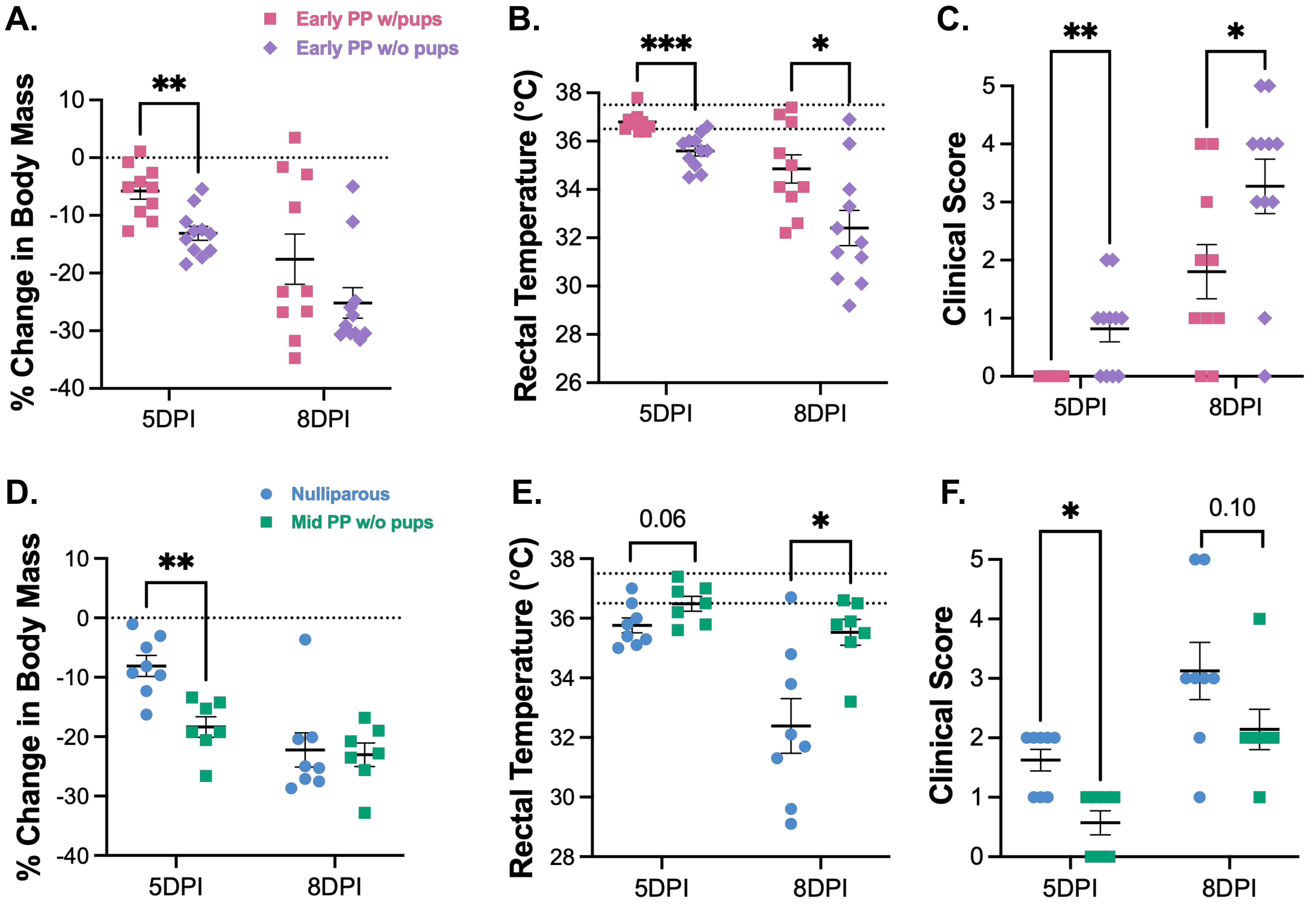

A.
Early PP w/pups
Early PP w/o pups
% Change in Body Mass
10
0
-10
-20
-30
-40
**
5DPI
8DPI
B.
Rectal Temperature (°C)
38
36
34
32
30
28
26
***
*
5DPI
8DPI
C.
Clinical Score
5
4
3
2
1
0
**
*
5DPI
8DPI
D.
Nulliparous
Mid PP w/o pups
% Change in Body Mass
10
0
-10
-20
-30
-40
**
5DPI
8DPI
E.
Rectal Temperature (°C)
38
36
34
32
30
28
0.06
*
5DPI
8DPI
F.
Clinical Score
5
4
3
2
1
0
*
0.10
5DPI
8DPI


Fig. S4.

**Fig. S4. The transition into lactation promotes resistance to severe disease following IAV infection.**

**(A-C)** Postpartum dams with and without pups were intranasally inoculated with a lethal dose of PR8 virus at 1-day postpartum. **(A)** Change in body mass, **(B)** core body temperature, and **(C)** clinical signs of disease are shown at 5- and 8-DPI (n = 10-11/group). **(D-F)** At 7-days postpartum, litters were removed and dams were intranasally inoculated with a lethal dose of PR8 virus with nulliparous females used as controls. **(D)** Changes in starting body mass, **(E)** core body temperature, and **(F)** clinical signs of disease are shown at 5- and 8-DPI (n = 7-8/group). For percent change in body mass, dashed lines at 0 indicate starting body mass. For rectal temperature, dashed lines indicate standard core body temperature of laboratory mouse (36.5 to 37.5 °C). Error bars represent the mean ± SEM from 2-4 independent experiments with individual mice indicated by symbols. $*P < 0.05$, $**P < 0.01$, $***P < 0.001$. Data were analyzed using multiple Unpaired t tests with Welch correction (A-B, D-E), and Mann-Whitney tests (C and F).

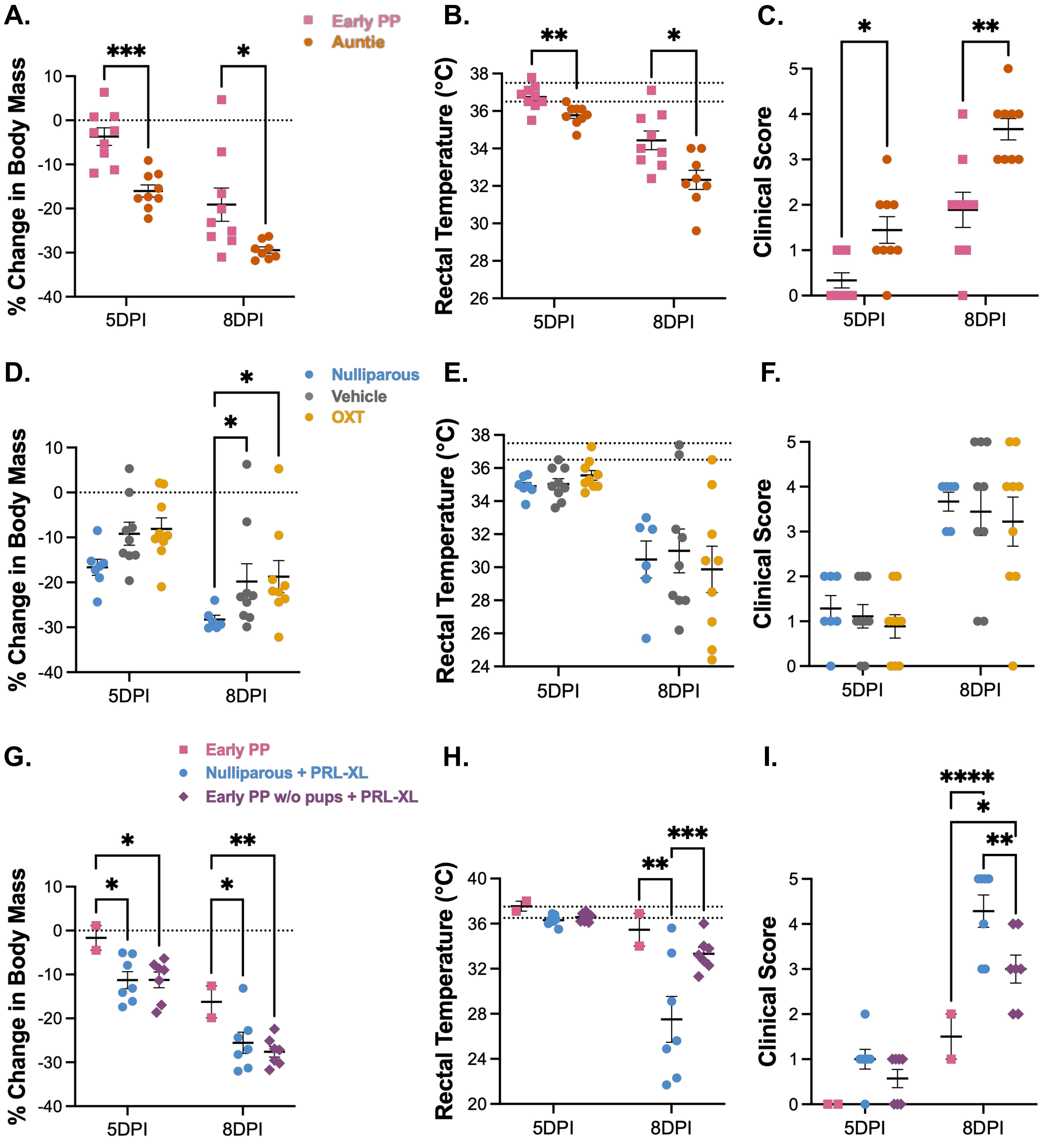

A.
Early PP
Auntie
% Change in Body Mass
***
*
5DPI
8DPI
B.
Rectal Temperature (°C)
**
*
C.
Clinical Score
*
**
D.
Nulliparous
Vehicle
OXT
E.
F.
G.
Early PP
Nulliparous + PRL-XL
Early PP w/o pups + PRL-XL
H.
I.
****
***


Fig. S5.

**Fig. S5. Integration of multiple lactation-associated signals are necessary for maternal resilience to IAV infection.**

**(A-C)** At gestational day 15.5, pregnant dams and nulliparous aunties were co-housed, followed by birth and subsequent intranasal inoculation with a lethal dose of PR8 virus at 1-day postpartum in both mice. **(A)** Change in starting body mass, **(B)** core body temperature, and **(C)** clinical signs of disease are shown at 5- and 8-DPI (n = 9/group). **(D-F)** For subcutaneous implantation of osmotic pumps with continuous delivery of oxytocin or sterile saline (vehicle), **(D)** the change in starting body mass, **(E)** core body temperature, and **(F)** clinical signs of disease are shown at 5- and 8-DPI (n = 7-9/group). **(G-I)** In prolactin-XL treated nulliparous females and postpartum dams without pups, **(G)** the change in starting body mass, **(H)** core body temperature, and **(I)** clinical signs of disease are shown at 5- and 8-DPI (n = 2-7/group). For percent change in body mass, dashed lines at 0 indicate starting body mass. For rectal temperature, dashed lines indicate standard core body temperature of laboratory mouse (36.5 to 37.5 °C). Error bars represent the mean ± SEM from 2-3 independent experiments with individual mice indicated by symbols. $^{*}P < 0.05$, $^{**}P < 0.01$, $^{***}P < 0.001$, $^{****}P < 0.0001$, ns $P > 0.05$. Data were analyzed using multiple unpaired t test with Welch correction (A-B), two-way ANOVA with Fishers LSD (D-F, G-I), and multiple Mann-Whitney tests (C).
